# Particle attachment drives seasonal abundance and photoheterotrophy of marine aerobic anoxygenic phototrophs

**DOI:** 10.1101/2025.04.22.649935

**Authors:** Cristian Villena-Alemany, Ana Vrdoljak Tomaš, Izabela Mujakić, Karel Kopejtka, Danijela Šantić, Michal Koblížek

## Abstract

**Background:** Aerobic Anoxygenic Phototrophic (AAP) bacteria are an essential component of aquatic microbial communities and play an important role in carbon cycling due to their ability to supplement their chemoorganotrophic metabolism with light-derived energy. While most of the previous studies focused on abundance, species composition and seasonal changes of AAP bacteria, their affinity for the particle-attachment did not attract much attention. Similarly, it remains unclear whether the entire AAP community is phototrophically active. This study investigated the seasonal changes in the composition of free-living and particle-attached AAP bacteria in the central Adriatic Sea’s coastal waters using both DNA and RNA *puf*M amplicon gene sequencing in the particle-attached and the free-living fractions.

**Results:** AAP bacterial abundance grew from 1.27 × 10^4^ cells mL^-1^ in winter to 8.30 × 10^4^ cells mL^-1^ in summer. The proportion of AAP bacteria was consistently higher in the particle-attached fraction, particularly in spring and summer. DNA and RNA *puf*M amplicon analyses revealed large differences in activity among the species forming the AAP communities. Additionally, DNA-based assessments underestimated the phototrophic activity of certain genera, demonstrating discrepancies between the gene presence and its functional activity.

**Conclusions:** Our data demonstrated that the expression of phototrophic genes in AAP bacteria is not uniform and largely varies throughout seasons and fractions. The particle-attached fraction harboured more than twice as many active AAP bacteria as the free-living fraction, with seasonal shifts and lifestyle driving changes in the phototrophy gene expression. RNA and DNA libraries revealed discrepancies between total and active AAP bacterial communities, emphasizing the necessity of transcript-based approaches for accurately assessing photoheterotrophic activity in marine environments. The pronounced partitioning of AAP bacterial diversity and activity between free-living and particle-attached fractions indicated the ecological specialization of certain AAP lineages, which may have noteworthy implications for the consumption of particulate organic matter and, ultimately, carbon cycling in coastal waters.

## Introduction

Bacteria perform fundamental roles in aquatic environments, driving key biogeochemical processes and shaping ecosystem dynamics [1]. Bacteria are typically divided as autotrophic or heterotrophic according to their carbon source, and they are critical for the primary and secondary production. There is an additional functional category that diverges from the aforementioned classification: the photoheterotrophic bacteria, which combine chemoorganotrophic metabolism with phototrophically harvested energy [2]. Photoheterotrophic bacteria use two fundamentally different systems for light harvesting: rhodopsin-pumps and bacteriochlorophyll (BChl) containing reaction centres, which may be combined in some species [3, 4]. Bacteria containing BChl*a*-containing RCs (e.g. purple bacteria) were previously associated to anoxic environments until aerobic anoxygenic photoheterotrophic (AAP) bacteria were shown to be a relevant part of the aerobic aquatic bacterial communities [5, 6].

AAP bacteria constitute a diverse metabolic group that is widely distributed across aquatic environments. Understanding their taxonomy, physiology and environmental patterns is essential for comprehending how their community operates. Their abundance and community composition exhibit strong seasonal changes [7–9] driven by primary producers and environmental phenology [10, 11]. Furthermore, cultures of AAP bacteria exposed to light reduced their respiration and accumulated more biomass [12–14].Infrared light, specially absorbed by Bchl*a* and therefore AAP bacteria, reduced the microbial respiration of natural planktonic communities and boosted the radiolabelled organic substrate uptake [15]. This effect has strong implication on the aquatic carbon cycle and its magnitude varies with the composition of the AAP community, showing that specific lineages may perform more photoheterotrophy, resulting in a larger impact.

In marine environments, the AAP communities displayed differential tendencies towards the free-living or particle-attached lifestyle [16]. Indeed, AAP bacteria generally correlated with dissolved organic carbon (DOC) and chlorophyll [17] and were more abundant in freshwater aggregates [18] accounting for up to 52% of the total bacteria in freshwater particle-attached communities [19]. Additionally, the taxonomy and ecology of overall microbial communities differed between free-living and particle-attached lifestyles [20–23] and, while AAP bacteria constitute important portion of marine bacterial community, little is known about their different community composition in particle-attached and free-living fractions.

The taxonomic composition of AAP bacterial communities is routinely analysed using the *puf*M gene, which encodes the M subunit of the anoxygenic type-II reaction centre [24–26]. Temporal and spatial analyses of the *puf*M gene documented that marine AAP bacteria represent a highly diverse and dynamic community [7, 27–30], which largely differs from the freshwater AAP species[8, 31–33]. Nonetheless the sole presence of the *puf*M gene did not guarantee the photoheterotrophic activity in freshwaters. Indeed, disparities between the presence of *puf*M genes and *puf*M transcripts from freshwaters documented that some AAP bacteria were more phototrophically active than what *puf*M gene libraries indicate [31]. Alternatively, the photoheterotrophic significance of relatively abundant *puf*M gene sequences might be overestimated if they are not actively expressed. Currently, it is uncertain whether the *puf*M gene amplicons in marine environments accurately reflect the active phototrophic community. Furthermore, the effect of free-living or particle-attached lifestyles on the phototrophic activity of the AAP communities remains unknown.

Here, we hypothesized that the composition of the phototrophically active (RNA) marine AAP community will differ from the overall (DNA) AAP community. Additionally, we expected that the particle-attached and free-living lifestyles will influence the AAP community composition and modulate the phototrophy gene expression patterns. To address the aforementioned questions, water samples from Adriatic Sea were collected in winter, spring and summer of 2023, and divided into two fractions containing a) free-living bacteria and b) particle-attached. Environmental factors, AAP abundance and biovolume, BChl*a* concentration and DNA and RNA *puf*M amplicon libraries were investigated.

## Materials and methods

### Sampling and environmental variables

Water samples were collected on 2^nd^ of February for winter, 5^th^ of May for spring and 7^th^ of July for summer 2023 from Kaštela Bay, in the eastern section of central Adriatic Sea (43°31’06.0"N 16°22’54.0"E). At 08:00, 50 litres were collected from 0.5 m depth and transported immediately to the laboratory. Ten litres were filtered through 150 µm mesh to remove larger plankton while preserving smaller particles (particle-attached fraction; PA) and subsequently, five litres were filtered through 1.2 µm pore size filter to remove particles and particle-attached bacteria (free-living fraction; FL). Between 350 and 1,000 mL of PA fraction and 650 – 2,000 mL of FL fraction were immediately filtered through a 47 mm of 0.45 µm pore size PES membrane filter (Millipore Express^®^ PLUS, Ireland), flash frozen in liquid nitrogen and stored at -80°C until nucleic acids extraction (at the end of the sampling). 0.45 µm pore size filter preferentially retain major proportion of AAP bacteria [16], which have larger average cell sizes than heterotrophic bacteria [34, 35]. Additionally, between 300 and 1700 mL were filtered through 25 mm of 0.45 µm PES membrane filter (Merk Millipore, Ireland) for BChl*a* quantification using HPLC, as described in Ruiz-Gazulla *et al*., 2024 [10]. Chlorophyll-*a* (Chl*a*) was determined from 500 mL subsamples filtered through Whatman GF/F glass fiber filters and stored at −20 °C. Filters were extracted in 90% acetone, and the extracts were analysed using a Turner TD -700 laboratory fluorimeter calibrated with pure Chl*a* (Sigma) [36] (File S1). AAP bacteria were counted using an epifluorescence microscope as described in Šantić *et al*., 2025 [37]. AAP bacteria biovolume of the PA and the FL fractions was measured in the microscope to confirm that 1.2 µm filtration had no effect on free-living AAP bacteria (File S2). The amount of reaction centres per cell was calculated as described in Ruiz-Gazulla et al., 2024 [10].

### RNA and DNA amplicon libraries

Total nucleic acids were extracted according to Griffiths *et al*., 2000 [38] with modifications [39], resuspended in 50 µL of DNase and RNase-free water (MP Biomedicals, Solon, OH, USA), and stored at –80°C. Both DNA and RNA amplicon libraries were generated using UniF-UniR primers [40] after demonstrating the highest coverage for marine AAP bacteria [26]. Prior to reverse transcription, DNA was depleted from total nucleic acids by digestion using the Turbo DNA-free kit (Ambion, Invitrogen) according to the manufacturer’s protocol. The absence of DNA in the RNA fraction was verified using PCR. cDNA was synthesized from 200 ng of total RNA using SuperScript IV VILO Mastermix (Invitrogen) according to manufacturer’s protocol. The obtained cDNA was diluted tenfold and used for the amplicon library generation by *puf*M gene PCR. RNA sample from particle-attached in spring could not be amplified. The samples were amplified in triplicate using Phusion HighFidelity PCR MasterMix (Thermo Scientific, USA) with the following PCR amplification protocol: initial denaturation for 3 min at 98°C, 30 cycles of 98°C for 10 s, 52°C for 30 s, 72°C for 30 s, and a final elongation at 72°C for 5 min. The triplicates were combined together and sequenced using Illumina MiSeq platform (2 × 250 bp) at Genomics Core Facility, Universitat Pompeu Fabra, Catalonia.

### Amplicon analyses

The primer sequences were removed using Cutadapts v1.16 with maximum error (-e 0.1 -n 1 -O 10) [41]. The initial number of reads ranged between 249,455 and 370,072 reads per sample (mean ± SD: 304,839 ± 36,669). The subsequent ASV construction was carried out in R/Bioconductor environment using DADA2 package v.1.29.0. Reads were truncated with *filterAndTrim*(truncLen=c(150,150)); forward and reverse reads were combined (*mergePairs()*); chimera were removed (*removeBimeraDenovo(method = "pooled"*) and 5,570 ASV (234,503±31,658 reads per sample) were constructed. Phyloseq file was generated in Phyloseq v1.45 for subsequent modifications [42]. Singletons, very rare ASVs (less than three reads in single samples) were removed as well as ASVs that poorly aligned using ClustalW v2.1 [43] in Geneious Prime v 2019.2.3 and verified not to be *puf*M genes using BLAST against the NCBI non-redundant database [44, 45]. The final dataset contained 1,143 *puf*M ASVs (230,386 ± 31,330 reads per sample, File S3). Samples were deposited in the NCBI database (BioSamples SAMN47815439 - SAMN47815449) under BioProject PRJNA1247470. The taxonomic assignment was performed using phylogenetic placement as described in Villena-Alemany *et al*., 2024 [11] (File S3). Plots were generated using ggplot2 v3.3.6 [46] in R v4.3.1 [47]. Shannon alpha diversity was calculated using *plot_richeness()* from Phyloseq v1.45. Beta diversity was assessed using principal component analysis on centred log ratio (CLR) transformed read counts using *transform()* from Microbiome v1.17.42. Genus pairwise comparison of the different fractions, seasons and amplicon libraries were obtained after aggregating the dataset to the genus level using *tax_glom()* from Phyloseq v1.45.

## Results

### Seasonal changes in Abundance of AAP bacteria and Bacteriochlorophyll-*a*

According to the BChl*a* concentration, the photoheterotrophy performed by AAP bacteria increased during the seasons, with the maximum concentration detected in the summer and spring (4.23 and 3.76 ng L^-1^, respectively, Figure 1A). BChl*a* concentration substantially contrasted between the particle-attached and the free-living fraction in the three seasons with the larger differences occurring in spring and summer, reaching more than five times in the particle-attached than in the free-living fraction in spring. Similar trend was observed in the AAP abundance, with the lowest numbers in winter (1.27 × 10^4^ cells mL^-1^) and the maximum abundance in summer (8.30 × 10^4^ cells mL^-1^, Figure 1A), constituting 13.8% of the total bacteria (File S1). Indeed, BChl*a* and AAP bacterial counts showed a significant (p-value=0.002) positive correlation (Figure 1B) allowing for the computation of 2.14 × 10^10^ - 8.51 × 10^11^ PSUs per litre and an average of 1.64 × 10^3^ - 1.12 × 10^4^ RCs per AAP cell.

**Figure 1.**
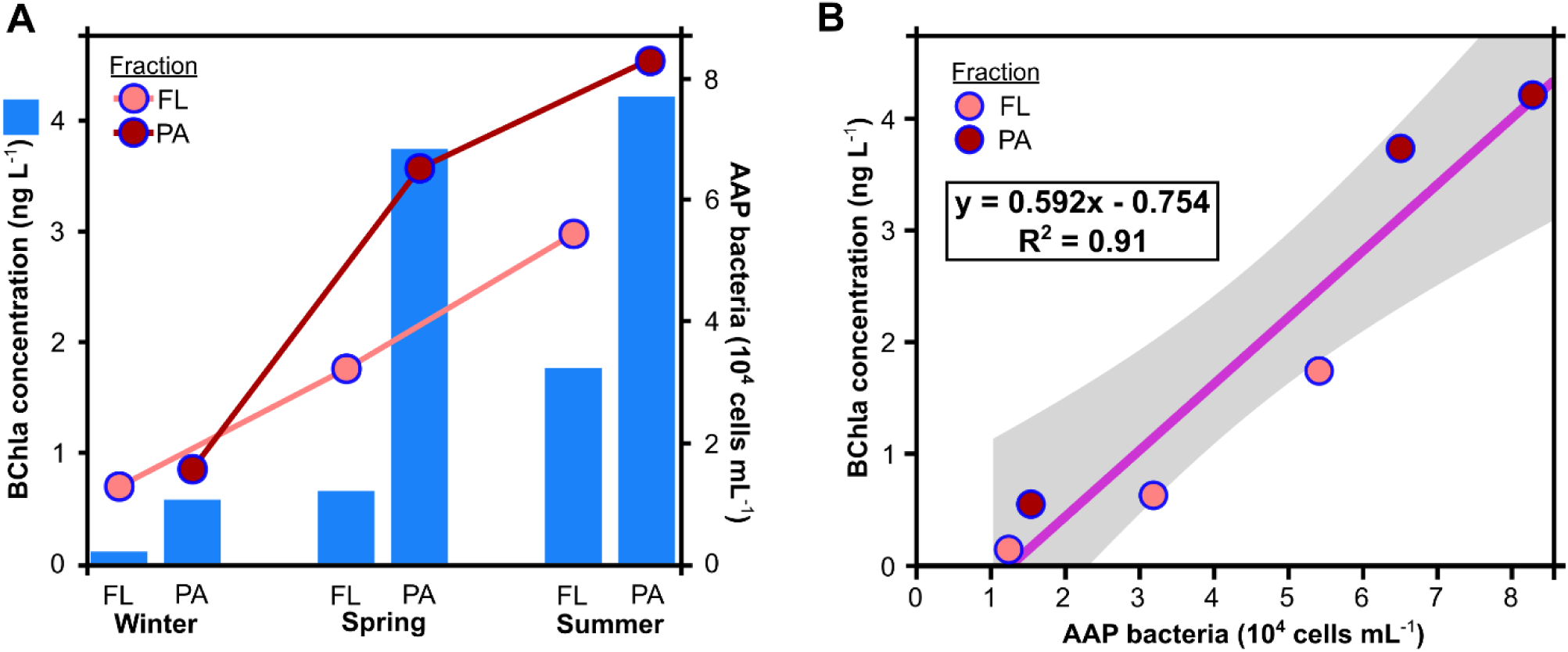
A) Bar plots of BChl*a* concentration and AAP bacteria abundance (lines) for the free-living (FL) and particle-attached (PA) fractions for the three seasons. B) Linear model of the correlation between AAP bacteria.

### AAP community composition

The beta diversity between the community composition of total and active AAP bacteria, based on DNA and RNA *puf*M gene amplicons, exhibited important differences at the seasonal level (Supplementary Figure 1). The total and active AAP community presented higher alpha diversity in winter and the lowest alpha diversity occurred in summer, particularly in the RNA library (File S3). In winter, the total AAP community was dominated by *Luminiphilus* while the active was more diverse (Figure 1). In spring, also *Luminiphilus* dominated in the total and the active community whereas in the summer, Arenicellales and Rhodobacterales HIMB11 (Figure 2). Overall, *Luminiphilus* contributed the most (21.1 – 58.6%) to the total AAP community throughout the year while its activity was highest in the spring and the lowest in the summer. Pairwise comparison of the percent contribution of each genus between seasons (Figure S2) revealed an internal turnover of Rhodobacterales genera, a reduction in Burkholderiales, and a rise in Pseudomonadales, mostly *Luminiphilus* in the transition from winter to spring. Alternatively, spring to summer transition of the whole AAP community indicated a similar dynamic succession of Rhodobacterales genera but, in both, presence and activity, there was a substantial increase in Arenicellales UBA668 and a drop in *Luminiphilus* contribution (Figure S2).

**Figure 2.**
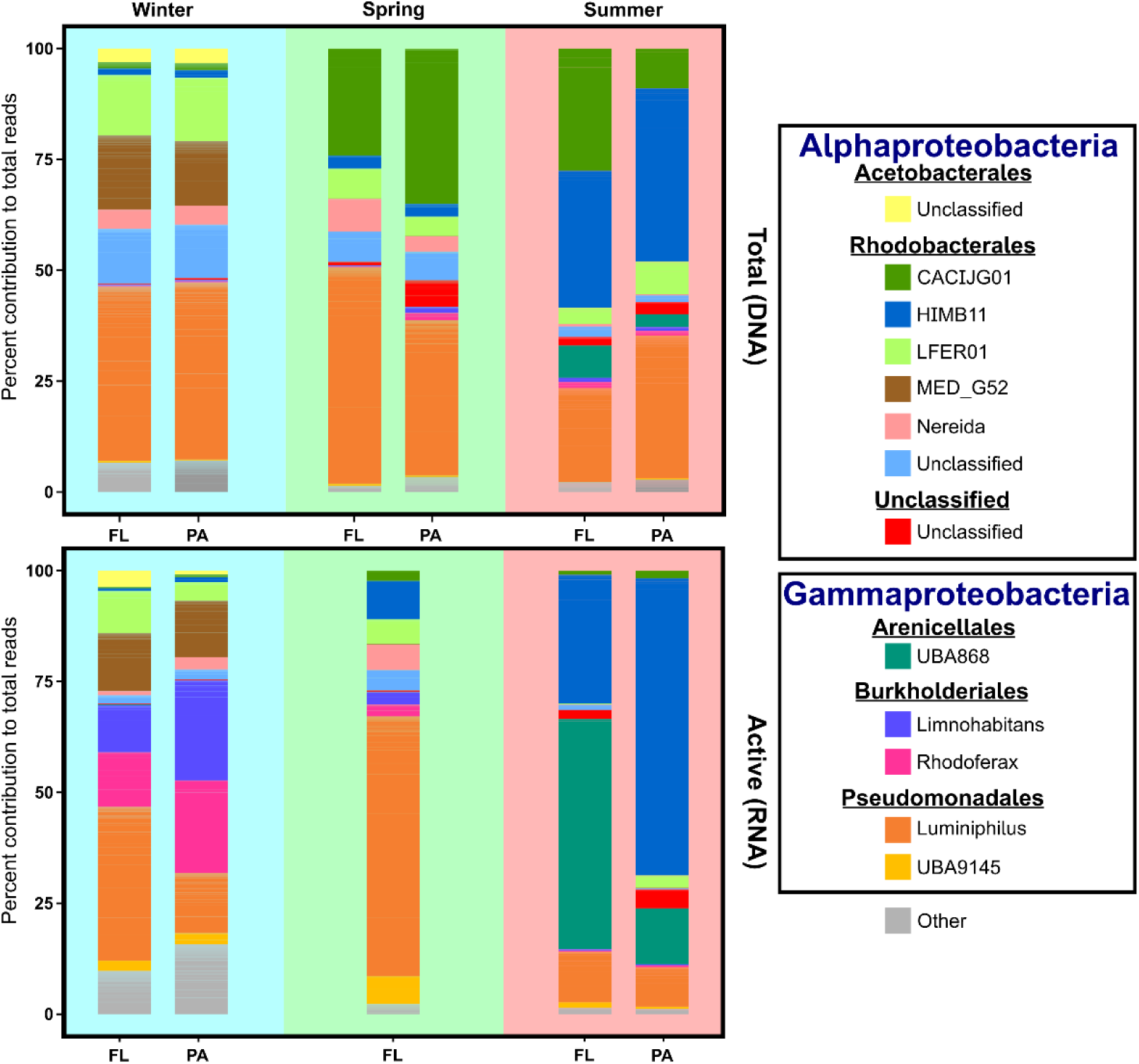
Percent contribution of AAP genera to community composition of the total (up) and of the active AAP community (down) for the free-living (FL) and the particle-attached with free-living (PA+FL). Genera less abundant than 0.5% are shown as “Other”.

### Particle-attached and free-living AAP bacteria

In concordance with the disparities in BChl*a* concentration and AAP bacteria abundance across fractions (Figure 1), the total AAP community and the expression of phototrophy genes also displayed differences between the FL and the PA fractions with magnitudes varying substantially between seasons (Figure 3). In winter, the AAP community composition was more evenly distributed throughout fractions. Nevertheless, *Luminiphilus* was more active in the FL fraction whereas Rhizobiales, *Limnohabitans*, *Rhodoferax* and other Burkholderiales tended to express their phototrophy genes when attached to particles. In spring there were larger differences between fractions in the total AAP community composition, with a repeated enrichment of *Luminiphilus* in the free-living fraction and a higher contribution of Rhodobacterales CACIJG01 in the particle-attached. In summer, the differences between fractions were larger in the active AAP community with a higher phototrophic activity of the free-living Arenicellales UBA868 and an important increase in Rhodobacterales HIMB11 *puf*M expression in the particle-attached fraction. *Luminiphilus* contributed more to the overall AAP community of the PA fraction but was slightly more phototrophically active when living freely during the summer.

**Figure 3.**
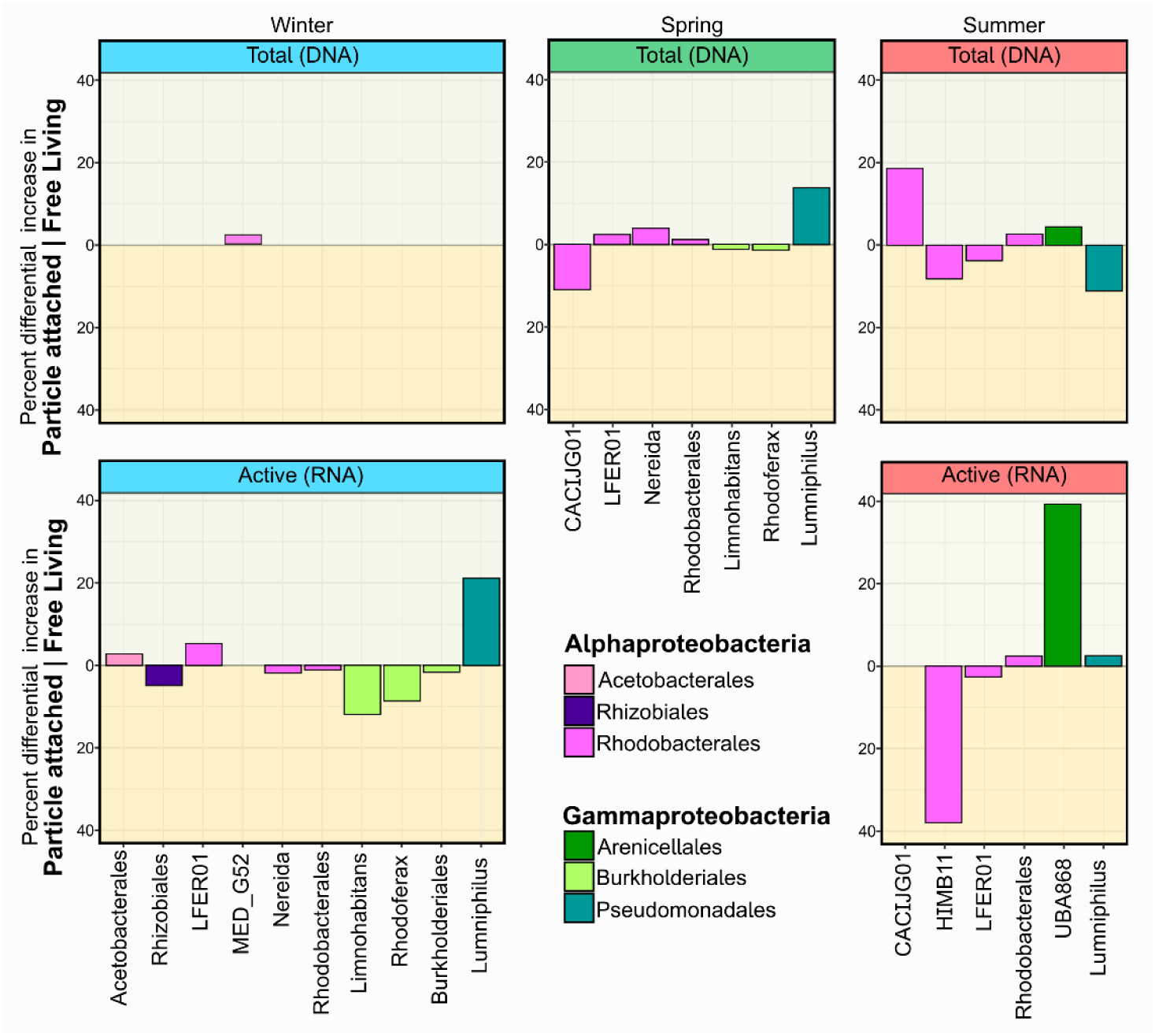
Pairwise comparison of the percent contribution of AAP genera between the free living and particle attached fractions for the total (up) and the phototrophically active (down) AAP community.

### Total and phototrophically active AAP community

Seasonal and fraction-dependent changes in community composition could be observed at both, RNA and DNA levels. Furthermore, there were important differences as well between the total and the phototrophically active AAP community (Figure 4). In winter, there was a similar general expression pattern in both fractions, with a large overrepresentation of Rhodobacterales and *Luminiphilus* in DNA libraries and an underestimating of the activity of several Burkholderiales genera (Figure 4). In spring, there was a heterogeneous behavior of Rhodobacterales genera and greater expression of the phototrophy genes, primarily *Luminiphilus* and UBA9115, but also *Limnohabitans* and *Rhodoferax*. In summer, the season with the highest AAP-derived photoheterotrophy, there was the largest difference between the total and the active AAP communities, with a large discordance in Arenicellales UBA868 contribution between fractions, largely overexpressing the phototrophy genes in the FL fraction in comparison to their contribution to DNA amplicon libraries. Rhodobacterales HIMB11 was underrepresented in the DNA libraries from the PA fraction and *Luminiphilus* was overrepresented in the DNA libraries.

**Figure 4.**
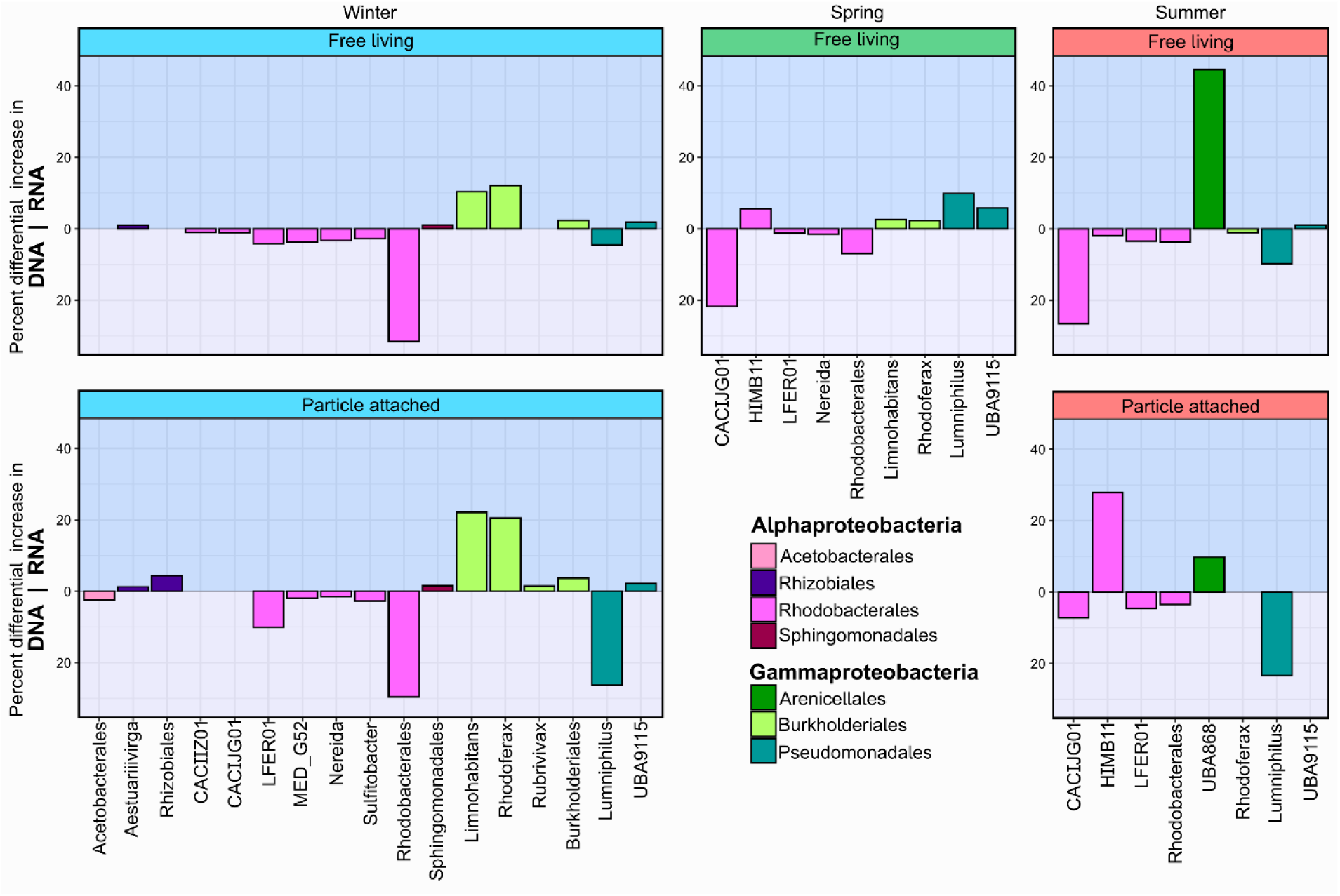
Pairwise comparison of the percent contribution of AAP genera to the RNA and DNA libraries for the free-living (up) and particle-attached (down).

## Discussion

AAP bacteria are found ubiquitously in aquatic environments. Due to their particular capability to supplement their metabolisms with light energy, they have been postulated to play a major role in the recycling of dissolved organic carbon, channeling it to upper trophic levels and substantially contributing to the microbial loop [48]. The abundance and diversity dynamics of AAP bacteria have been investigated in diverse marine habitats [7, 27, 49, 50], including the Adriatic Sea [28, 51, 52]. Their abundance, usually ranged between 0.1 and 11% of the total bacterial community as shown in North Pacific [53], Baltic Sea [9, 54], Artic Ocean [55], Mediterranean Sea [56–58] and in the Adriatic Sea [28, 51]. The community composition of AAP bacteria in marine environments, has commonly been addressed using amplicon analysis of the *puf*M gene. Nevertheless, the presence of the genes did not guarantee their expression in freshwaters [31] and, DNA amplicon libraries may largely differ from RNA amplicon libraries in marine environments as well.

The lifestyle preference of AAP bacteria towards an attachment to particles has been suggested both, in marine and freshwater environments [16, 59]. It has been shown their affinity for carbon rich sources, which may explain their declination toward a particle-attached lifestyle [10, 11, 60]. Additionally, AAP bacteria thrive during periods of high primary production and accounted for 20% of the FL bacteria and up to 52% of the PA bacteria in freshwaters under that circumstances [18, 19]. Nevertheless, the preference of AAP bacteria towards the PA or the FL lifestyle in marine environments remained unknown. Furthermore, AAP bacteria is a highly diverse metabolic group with various phylogenetic and ecological intragroups, and whether certain AAP bacteria prefer to live attached to particles or free-living, remains undisclosed. Finally, little is known about how the phototrophic activity of AAP bacteria varies over the fractions and seasons.

In the current study, the AAP contribution to the total bacterial abundance was larger (up to 13.8%, File S1) than the average reported in open ocean and Mediterranean Sea and in concordance with prior studies from the Adriatic Sea [28, 53, 56, 61] (Figure 1A). AAP bacteria have shown to prefer more productive coastal waters [62]. Additionally, there was a marked seasonality in the abundance and phototrophy of AAP bacteria with higher concentration of AAP bacteria and BChl*a* in more productive seasons (productivity according to the prokaryotic primary producers, File S1). In contrast to previous observations [28], the highest AAP abundance occurred in summer and both, the AAP abundance and phototrophy by AAP bacteria were the maxima reported in this area [28, 51, 61] but in concordance with the concentrations observed in other marine environments [27].

Furthermore, the direct relationship between the AAP abundance and BChl*a* (Figure 1B) indicated that AAP bacteria had an overall homogenous number of reaction centers across the season and the fractions. As previously proposed [10], we here support that is plausible, assuming a constant number of BChl*a* molecules per PSU, a comparable amount of BChl*a* molecules per AAP bacteria.

DNA and RNA *pufM* gene amplicons reflected the seasonal variability in the total and in the phototrophically active AAP community composition, but the biggest differences were detected in the RNA libraries (Figure 2). Similarly, the variation in the lifestyle preferences of AAP bacteria could be observed to a greater extent, in the RNA libraries. As previously documented for freshwater [31], the potential and the active AAP community differed and RNA amplicon libraries better provided a snapshot of the AAP phototrophic activity and its implications in the environmental functioning. Additionally, the greater discrepancies between fractions of RNA and DNA libraries pointed out that specific AAP bacteria tend to express their phototrophy genes depending whether they are particle-attached or free-living. This finding suggests that expression of phototrophy genes is regulated, in addition to the carbon availability observed in cultures [63] and to light in natural environments [31, 49, 64], by whether they are-attached to particles or live freely. Yet it is true that the nature of the particles can vary greatly and we did not cover their composition. Therefore, whether the responses of AAP bacteria to the different lifestyle could be impacted by the nutrient composition of the particles will remain, so far, undisclosed.

Furthermore, there were disparities between the presence of specific AAP bacteria and their phototrophic activity among seasons and fractions. For instance, *Luminiphilus*, a previously reported dominant AAP bacteria in central Adriatic Sea [28], came out to be the most common in winter and spring, but presented lower contribution to the AAP community in summer. The discrepancies between the community composition of previous work in the same Kaštela Bay is partially explained by the use of different *puf*M primers. Here, primers were chosen after extensive coverage analysis [26] that indicated UniF-UniR primer pair [40] to contain the best coverage in marine environments. Indeed, the RNA libraries pointed out that the presence of *Luminiphilus* in both fractions based on DNA libreries is largely overestimating their role in the phototrophy of AAP community (Figure 2, Supplementary Figure 2). Additionally, the discrepancies between their high contribution in the PA fraction and their high activity in the free-living fraction in summer correspond with the potential the downregulation of phototrophy genes in a rich carbon environment that some AAP bacteria have shown [65].

An interesting observation is the occurrence of transitional or potential freshwater AAP bacteria such as Rhizobiales, *Limnohabitans* and *Rhodoferax* in winter and spring samples (Figure 3) as a result of the fresh water input from nearby river Jadro and the rain runoff. These sequences, were not just present consequence of an allochthonous transportation but, remarkably, they were active, especially in the particle-attached bacteria (Figures 3 and 4). In concordance, Arenicellales UBA868, a genus previously reported from estuary [66] and HIMB11 from coastal waters [67] were highly active in free-living and particle-attached, respectively, and confirmed the transitional nature of the Kaštela Bay [61]. Indeed. HIM11 represented more than 66% of the active AAP bacteria community when the AAP abundance and their phototrophy was the highest, in summer. Additionally, Rhodobacterales CACIJG01, previously reported from the Mediterranean Sea [68], highly contributed to the total AAP community but was clearly overrepresented when accounting for their activity. Finally, it is necessary to stress that both DNA and RNA libraries were done from the morning environmental communities and the expression of phototrophy genes have shown diel changes, not just in the abundance of their transcripts but also in the composition [31]. Therefore, phylotypes contributing importantly to the AAP community that here are shown as phototrophically inactive might have different patterns of gene expression than the overall AAP bacteria, as is the case for those one from the phylum Gemmatimonadota [69]. For instance, diel high resolution transcriptomics analysis of AAP bacteria would better confirm whether the expression patterns of phototrophy genes in marine AAP bacteria are daily downregulated by light and provide understanding of the photoheterotrophy by AAP bacteria in marine environments and its potential impacts in the microbial community and the carbon cycle.

## Conclusions

Active AAP bacteria have shown to contribute in absolute and relative numbers to a large extend of the bacterial composition during spring and summer. There was higher occurrence of AAP bacteria in the particle-attached fraction, and the expression of phototrophy genes was conditioned by their lifestyle. Furthermore, we confirmed that *puf*M DNA libraries draw distorted patterns of the active AAP community composition in marine environments and documented, mostly in winter, the phototrophic activity of phylotypes classically associated with freshwaters. Finally, we have shown that AAP community is a highly heterogeneous metabolic group and each genus presented inherent niche preferences to live and to be phototrophically active.

## Ethics approval and consent to participate

Not applicable

## Consent for publication

Not applicable

## Availability of data and materials

The dataset supporting the results and conclusions are embedded in the manuscript and available in the Sequence Read Archive (SRA) under Biosamples SAMN47815439 - SAMN47815449 from Bioproject PRJNA1247470.

## Author’s contributions

CVA and MK did the conceptualization; CVA, IM; formal analysis was carried out by CVA; investigation and experimental processes were done by CVA, AVT, IM, KK and DS; CVA wrote original draft and all the authors helped during the review and editing of the manuscript; CVA, AVT, IM and DS performed the sampling and measured the environmental variables. All authors have revised and approved the submitted version.

## Acknowledgements

The authors want to thank Ante Marasović, Slaven Jozić and Iva Stojan for help with sampling.

## Supplementary Figures

Figure S1: Principal component analysis of centered log-ratio transformed AAP community. Each dot represents a DNA (circle) or RNA (triangle) sample. The colour indicates the season: blue for winter, red for spring and green for summer.

Figure S2: Pairwise comparison of the percent contribution of AAP genera between transition of winter to spring (up) and the transition spring to summer (down) for the DNA and RNA libraries in the free-living and particle-attached.

## Supplementary Files

File S1: Environmental variables including physical and chemical parameters as well as biologic measurements.

File S2: AAP biovolume values per season and per fraction including number of counted cells, average and standard deviation of the biovolume in µm^3^. Additional ANOVA analysis that confirmed the no difference of biovolume between fractions.

File S3: Taxonomy of the ASVs (Taxonomy); reference ASVs identification and their sequence and ASV table including the ID of the sample, the Shannon alpha diversity index per sample and the number of reads associated to each ASV in each sample.

