## Supplementary figures and images for "Particle attachment drives seasonal abundance and photoheterotrophy of marine aerobic anoxygenic phototrophs"

### Figure S1

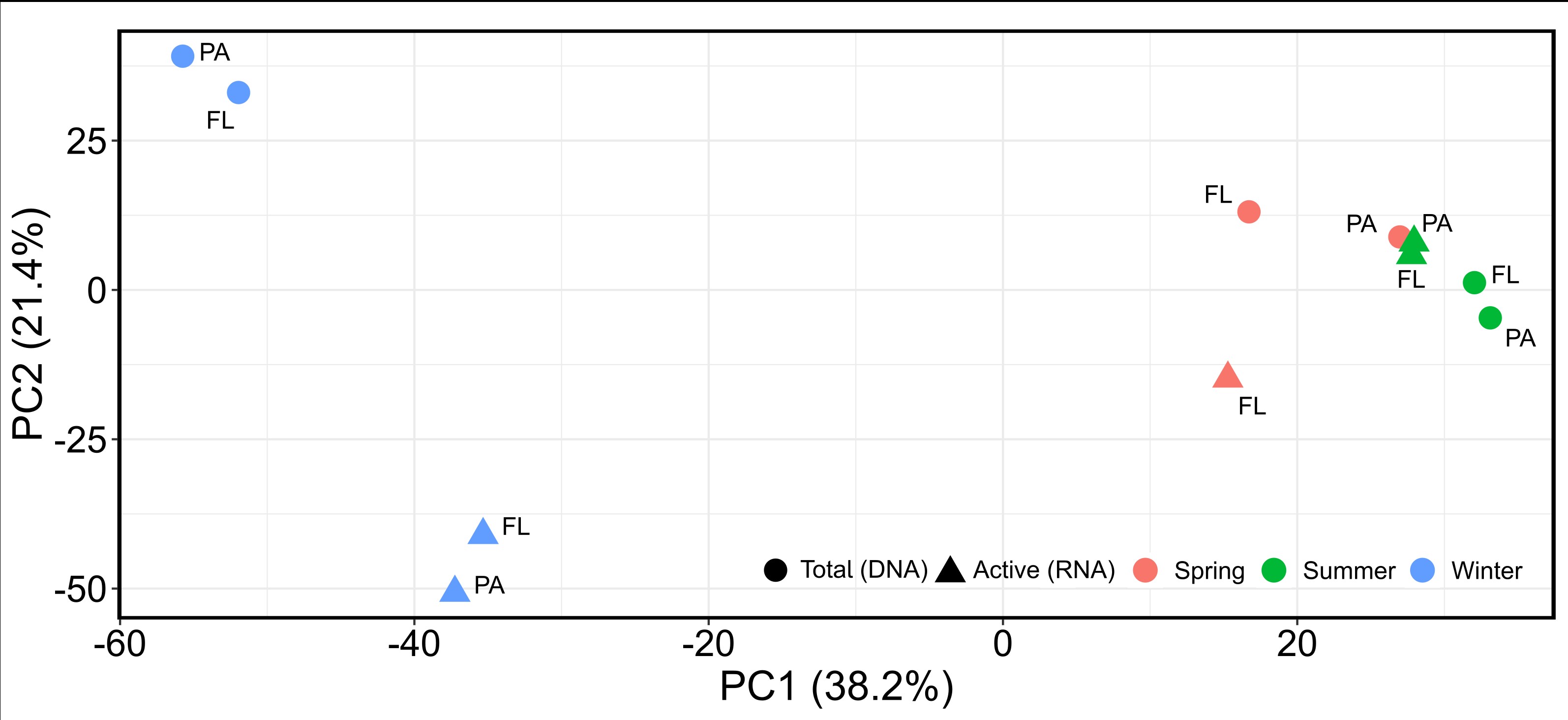

### Figure S2

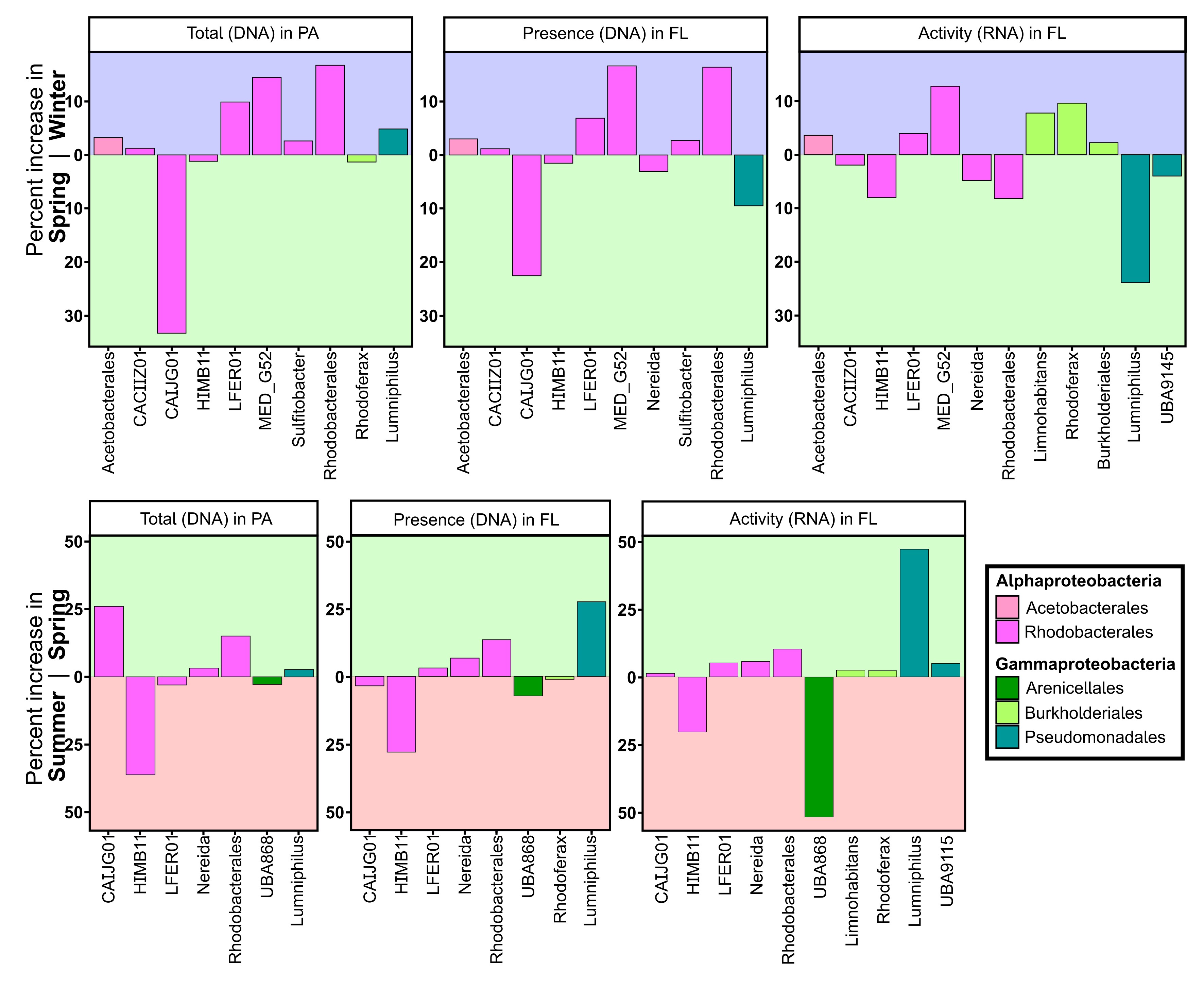
